# Balancing Lucidity: Muscle and Vestibular Stimulation for Lucid Dream Induction

**DOI:** 10.64898/2026.03.11.711028

**Authors:** Emma Peters, Xinlin Wang, Kathrin Fischer, Nicole Bühler, Nicolas Morath, Johanna Heitmann, Eva Nussbaumer, Ralf Kredel, Simon Maurer, Martin Dresler, Daniel Erlacher

**Author notes:** Corresponding Author: Emma Peters.

## Abstract

Lucid dream (LD) induction using external sensory stimulation has most commonly relied on distal cues such as lights or auditory signals, with mixed success rates. In this study, we investigated whether more direct bodily stimulation targeting the muscle and vestibular systems could influence LD induction. We compared electrical muscle stimulation (EMS) and galvanic vestibular stimulation (GVS), each combined with a two-week cognitive training protocol including dream journaling, reality checks, and association training. Twenty-eight participants (14 per group) completed two counterbalanced morning naps: one with stimulation (STIM) during REM sleep and with one sham-stimulation control (SHAM). EMS and GVS stimulation did not lead to increased incorporation of the stimulus. Lucidity rates were high in both EMS conditions, highlighting the substantial role of elevated baseline lucidity in induction studies, cognitive training, and expectation effects. In contrast, GVS stimulation significantly increased externally rated lucidity and DLQ questionnaire scores compared to control. Overall, the findings indicate that galvanic vestibular stimulation can increase dream lucidity. Future work should further examine the mechanisms by which vestibular stimulation influences dream awareness and its potential role in lucid dream induction.

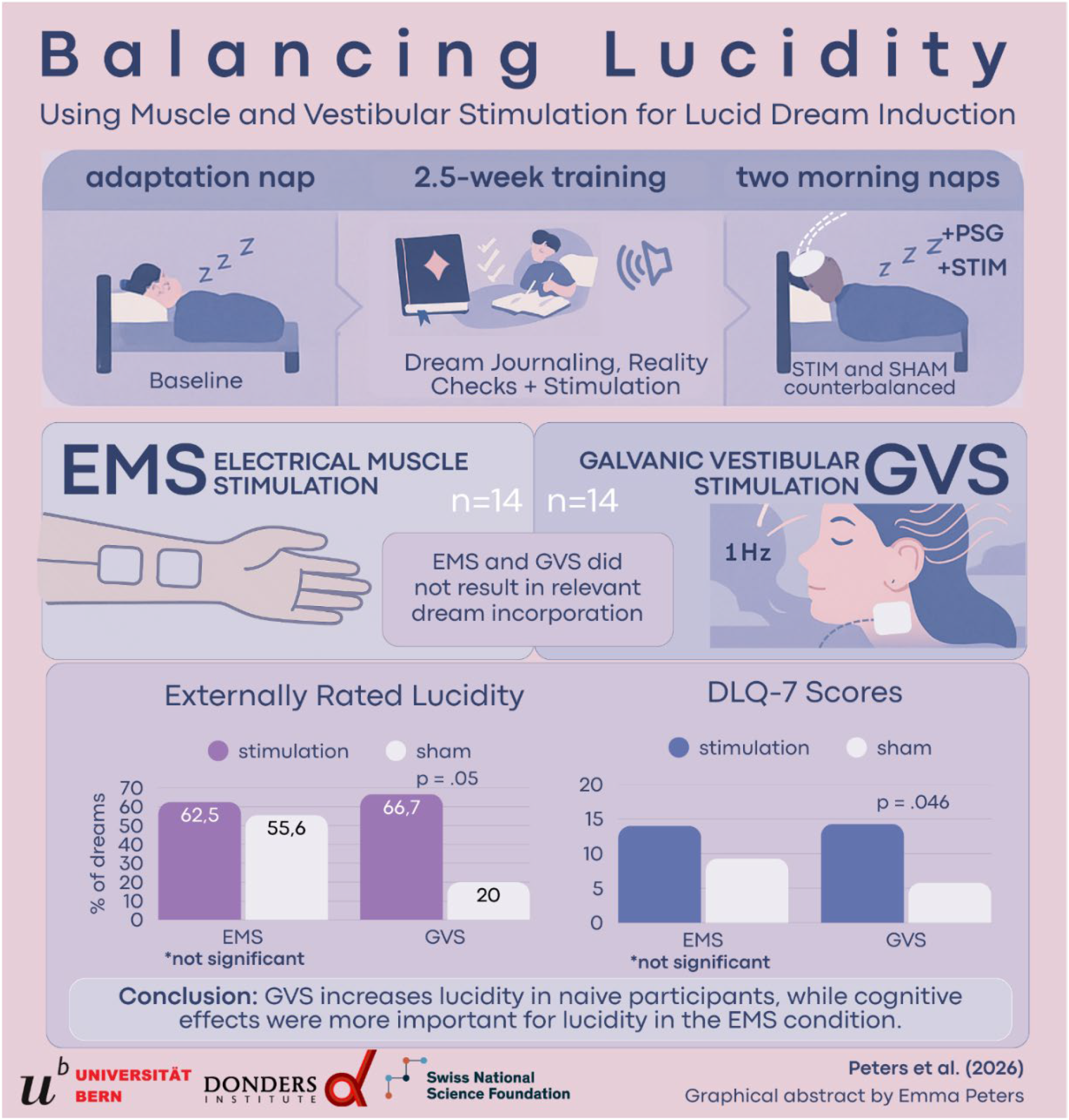

## Introduction

Lucid dreaming is a hybrid state of consciousness in which individuals become aware that they are dreaming while remaining immersed in the dream experience (Baird et al., 2019). During lucid dreams, dreamers may gain partial control over their actions and the dream environment (LaBerge & Ornstein, 1985). Although spontaneous lucid dreaming is relatively rare, it has attracted growing scientific interest due to its potential applications in nightmare treatment (Ouchene et al., 2023; Tzioridou et al., 2025), creativity and problem solving (Stumbrys & Daniels, 2010), and motor skill learning (Bonamino et al., 2023; Peters et al., 2023). Dream experiences are shaped not only by memory, emotion, and imagination, but also by signals originating from the sleeping body. External and internal sensory information can be incorporated into dreams, a phenomenon known as dream incorporation (Salvesen et al., 2024; Solomonova & Carr, 2019). Previous research has demonstrated that sensory cues presented during sleep, such as auditory tones, light flashes, or tactile stimulation can be integrated into dream content either directly or indirectly (Salvesen et al., 2024). When paired with presleep cognitive training, such cues may facilitate lucid dream induction through approaches such as Targeted Lucidity Reactivation (TLR; Carr et al., 2023).

Most lucid dream induction studies have relied on distal sensory cues, particularly visual and auditory stimulation (Tan & Fan, 2022). However, bodily sensations are central to dream experience, especially in lucid dreams, where dreamers often report heightened awareness of movement, balance, and bodily agency (Erlacher & Schredl, 2008). Dreamed movements engage sensorimotor brain regions in a manner comparable to executed movements, suggesting a close neurophysiological overlap between the physical and dreamed body (Dresler et al., 2011). These findings raise the possibility that stimulation targeting bodily systems more directly may provide effective cues for influencing dream awareness.

One such approach is electrical muscle stimulation (EMS), which induces brief muscle contractions and somatosensory feedback. Despite REM-related muscle atonia, EMS-induced sensations can reach the dreaming brain and be incorporated into dream content (Krugliakova et al., 2025; Peters et al., 2024; Salvesen et al., 2024). Tactile and proprioceptive stimulation has previously been associated with isolated cases of signal-verified lucid dreams (Paul et al., 2014), but systematic evaluations of EMS for lucid dream induction under controlled conditions remain limited. Another bodily system closely linked to dream phenomenology is the vestibular system. Vestibular sensations such as flying, floating, or falling are frequently reported in lucid dreams and have been associated with increased dream body awareness and control (Barrett, 1991; Picard-Deland et al., 2020). The vestibular system contributes to bodily self-consciousness, including self-location and self-agency, which are also present during dreaming (Picard-Deland et al., 2022). Galvanic vestibular stimulation (GVS) offers a non-invasive method to activate vestibular afferents during sleep (Krugliakova et al., 2025), yet its potential role in dreaming remains largely unexplored. Although both somatosensory and vestibular stimulation have been shown to influence dream content, it remains unclear whether these modalities differ in their ability to facilitate lucid dreaming, particularly when combined with structured cognitive training and compared against no-stimulation control conditions. Moreover, it is unknown whether direct incorporation of a bodily stimulus is necessary for enhancing lucidity, or whether more indirect modulation of dream phenomenology may be sufficient.

The present study addresses these gaps by testing EMS and GVS within a controlled laboratory design. Each stimulation modality was combined with a cognitive training protocol and evaluated using a within-modality comparison between stimulation (STIM) and no-stimulation (SHAM) conditions during REM sleep. Lucidity was assessed using dream report analysis and subjective questionnaires.

## Methods

### Participants and Ethics

Participants (n=28) were recruited through advertisements in social media groups and on the campus of the University of Bern, Switzerland. Exclusion criteria included low dream recall (<3 dreams/week), late bedtimes (>01:00), prolonged sleep latency (>1 h), a personal or family history of epilepsy, sleep-related or current psychiatric disorders, or a lucid dream frequency greater than once per month. The Faculty of Human Sciences ethically approved the study. All participants provided written informed consent and received financial compensation. Data protection was ensured through pseudonymous coding.

### Design and Procedure

Participants were assigned to the EMS or GVS group using alternating assignment at enrollment to maintain balanced group sizes. The study followed a 2.5 to 3-week protocol comprising an adaptation nap, a training phase, and two counterbalanced test naps (one with stimulation, one with sham stimulation). Each participant completed three morning nap sessions in total. An adaptation nap (day 1) was conducted without polysomnography (PSG) and sensory stimulation. Participants arrived at 8:00 AM and gave informed consent before the nap. They slept until 11:30 AM and were excluded if they reported significant sleep difficulties or no dream recall. If they were excluded at this point, another participant was recruited to reach the required sample size. The training phase (week 2) occurred between the adaptation and test naps, with participants completing a cognitive training protocol consisting of daily dream journaling, reality checks, and six 10-minute association training sessions, scheduled flexibly across afternoons. The test naps (week 3) were conducted on two consecutive mornings, with sessions starting at 7:00 AM. Each participant received one stimulation (STIM) nap (EMS or GVS) and one sham stimulation (SHAM) nap, with the order counterbalanced across and within groups. The pre-sleep audio protocol, adapted from Carr et al. (2023), was administered before sleep.

### Polysomnography

PSG, including electroencephalography (EEG), electrooculography (EOG), and electromyography (EMG), was utilized to monitor sleep stages based on AASM guidelines (Iber, 2007) during the test naps. PSG setup adhered to the 10-20 system, utilizing 32 active gLADYBIRD electrodes with the gUSBamp EEG system (Klem et al., 1999), using two horizontal EOG, chin EMG, and right earlobe reference. The sampling rate was 256 Hz. Experiment management and streaming were done using the University of Bern’s custom experiment management system Streamix. The sleep room had speakers, a microphone, and a camera for further observation and communication. PSG was administered solely to monitor sleep stages, not for any analytical purposes.

### Stimulation Methods

The study tested two different stimuli: Galvanic Vestibular Stimulation (GVS) and Electrical Muscle Stimulation (EMS), with a stimulus duration of two seconds.

### Galvanic Vestibular Stimulation (GVS)

Binaural GVS was administered using the NeuroConn DC-Stimulator Plus (*NeuroConn DC-STIMULATOR PLUS* | *Neurocare Technology*., 2023), and two 4.5x3.5mm carbon rubber electrode pads behind the ear over the mastoid bone. The electrodes were cut to a size that enabled precise application on the mastoid bone. Based on previous research, the stimulus parameters were set to a level where the participants felt a small sway sensation without feeling an uncomfortable skin sensation from the current behind the ear while sitting on the bed with their eyes open (intensity: varying from 1mA to 3mA peak-to-peak, frequency: 1Hz, duration: 2s, phase:0*, fade in/out: 0s, number of cycles: 2*2pi) . These parameters resulted in a quick left-right-left-right head sway. The amplitude varied across participants from 1000 µA to 2500 µA and was individually calibrated to elicit a clear vestibular sensation without discomfort.

### Electrical Muscle Stimulation (EMS)

Two 50x100mm self-adhesive TENS-EMS gel electrodes were placed on the volar side of the left forearm, targeting the flexor muscles responsible for ring-finger flexion. The distal electrode was positioned approximately 2–3 cm proximal to the wrist crease, aligned longitudinally with the forearm on the ulnar side. The proximal electrode was placed 5– 6 cm above the distal electrode along the same axis. This placement corresponded to the anatomical course of the wrist flexor muscle group involved in flexion of the ring finger. The Rehamove 3 “ScienceMode” Device facilitated electrical stimulation, allowing external trigger control via connection to a computer (*RehaMove— Funktionelle Elektrostimulation* | *HASOMED GmbH*., 2023). Before sleep, stimulation intensity was calibrated while participants were awake, using the lowest current that induced visible ring and pinky finger flexion without discomfort. EMS was delivered with a pulse period of 20ms (corresponding to 50 Hz), no amplitude ramp (ramp = 0), and a waveform defined by a single stimulation point of 3 mA with a pulse duration of 500 µs; the stimulation was applied as a continuous pulse train lasting approximately 2 seconds.

### Association training protocol

Participants completed daily dream journaling and six guided association sessions across the training period. During each 10-minute session, a recorded audio track was played while the 2-second EMS or GVS stimulation was administered with an interstimulus interval of 60 seconds.

### Targeted Lucidity Reactivation Protocol

The presleep protocol, adapted from Carr et al. (2023), was administered to reactivate the trained association between stimulation and lucidity. Participants were reminded of their task to become lucid and perform an LRLR eye-movement signal while attending to the stimulation. The protocol comprised 5 minutes of stimulation at 1-minute intervals paired with verbal lucidity prompts, followed by 8 minutes of stimulation at 1-minute intervals without verbal cues, and three final stimulations spaced 2, 4, and 7 minutes apart. After the final stimulation, participants were instructed to fall asleep, which marked the Lights Out time point.

### REM stimulation Protocol

REM periods were identified online from the PSG recordings using AASM criteria. REM sleep onset was defined as the first 30-second epoch showing low-amplitude mixed-frequency EEG activity, rapid eye movements in the EOG channels, and reduced chin EMG tone. Stimulation was applied only during clearly identifiable and stable REM periods. After REM onset, a 5-minute timer was started. Following the 5-minute wait period, a series of 2-second stimulations at a 30-second interstimulus interval (ISI) was presented continuously during REM sleep, as identified online via PSG monitoring. A maximum of 30 stimulations was presented. In the control condition, sham stimulation was applied by using a second timer that matched the STIM condition in time. If a scheduled REM period did not occur or was interrupted by wakefulness, the corresponding STIM or SHAM trial was postponed to the next available REM period rather than excluded. Consequently, participants did not always complete the same number of trials, and the assignment of conditions to specific REM periods varied across individuals. Nevertheless, the predefined counterbalancing ensured that, at the group level, STIM and SHAM trials were distributed across REM periods with similar frequency.

### REM-awakenings

REM awakenings were initiated after a participant woke up naturally or by calling the participant by name and asking whether they were awake. Once the participant confirmed being awake, they were asked: ‘What went through your head before you woke up?’ After the participant completed their report, the follow-up question ‘Did you notice anything else?’ was asked to gather additional details. Participants completed the Dream Lucidity Questionnaire (DLQ-7) (Stumbrys et al., 2013). Dream reports were audio-recorded. If time remained in the nap, participants were allowed to return to sleep, and the stimulation protocol was reinitiated during the next REM phase.

### Data Analysis

Audio dream reports were transcribed into written form and processed according to the Dream Report Conversion Manual (Schredl, 2010), removing non-dream-related content. Because the dream reports were collected in several languages, all reports were translated into English using DeepL Translator to ensure high-quality and contextually appropriate translations. The translated reports were then randomized and evaluated by two external raters who were blind to the study conditions.

### Dream Incorporation

Dream incorporation of the presented stimuli was analyzed following the methods used by Peters et al. (2024c). Two independent researchers scored all items. Inter-rater agreement was 89.5%. and disagreements were resolved through consensus scoring by a third researcher. For both modalities, direct mentioning of the stimulus was scored. Additionally, for GVS, the scale: ‘Balance-Related Activities’ was scored using the question: *‘Does the dreamer perform activities involving balancing and balance, e*.*g*., *flying or going over a narrow bridge? Does the dreamer have problems with balance, e*.*g*., *the occurrence of dizziness, fear of looking deep, etc?’*. For EMS, the two scales: ‘Movement of the Arms’ (Movement) and ‘Tactile or Somatosensory Sensations of the Arms’ (Tactile) were used with the questions: *‘Does the dreamer perform activities that have to do with the arms, e*.*g*., *waving, manual labor, boxing? The activity should be explicitly named*.*’* And *‘Does the dreamer experience activities in which a tactile or somatosensory sensation/stimulus plays a role?’*

### Lucidity Measurements

Three measures of lucidity were used to assess the prevalence of lucid dreaming in this study:

1. *External Rating of Lucidity (Ext_LD):* If the dream report showed any signs of explicit dream awareness or signs of metacognitive thinking or deliberate intention, Ext_LD was scored as present.
2. *Dream Lucidity Questionnaire (DLQ-7):* This questionnaire is designed to evaluate dream awareness (e.g., knowing one is dreaming, recognizing the unreality of the dream), control (e.g., ability to make choices, alter dream content, defy physical laws) and recall (e.g., remembering waking intentions or real-life facts). The DLQ includes 12 items, each rated on a 5-point Likert scale from 0 (‘not at all’) to 4 (‘very much’). In our study, we excluded item 7, as this item is seen as not reflective of lucid insight.

### Statistics

A generalized linear mixed model with a binomial distribution and logit link was conducted using Jamovi (The jamovi project, 2025) to test for differences in lucidity and dream incorporation under different stimulation conditions. The dependent variables were *Balance, Movement of the Arms, Tactile or Somatosensory Sensations in the Arms*, and *Ext_LD*. Participant was included as a random intercept. A linear mixed model was conducted to examine whether stimulation affected dream lucidity scores (DLQ-7), with participant included as a random intercept. Externally rated lucidity (binary outcome) was analysed using a generalized linear mixed model (GLMM), whereas DLQ-7 scores (continuous outcome) were analysed using a linear mixed model. Because these analyses targeted a small number of predefined and conceptually related lucidity outcomes and were evaluated using different models appropriate to the data structure, no formal correction for multiple comparisons was applied.

## Results

### Participants

A total of 28 participants were included in the study, with 14 participants per group. Group characteristics, including sex distribution, age, dream recall frequency, and lucid dream history, are summarized in Table 1.

### Awakenings and Dream reports

Across 28 participants, 12 reported at least one dream in both the STIM and SHAM conditions, 12 recalled a dream in only one condition, and four could not recall any dreams and were excluded from further analyses. In the EMS group, 13 participants were included. In the STIM condition, 13 dreams were included, and in SHAM, 16 dreams were included. In the GVS group, 11 participants were included. 12 dreams were included in STIM and 15 dreams in SHAM. The number of dreams per participant varied; some reported multiple dreams, and others none. Thus, one dream per participant per condition was included in the analysis by selecting the dream that contained an externally rated lucid dream (ExtLD) and/or the highest LDQ-7 score. This selection approach was chosen to maximize sensitivity to LD. Following this selection process, the final analysis included 8 EMS-STIM and 9 EMS-SHAM dreams, and 9 GVS-STIM and 10 GVS-SHAM dreams.

**Table 1.** Participant Overview.

| Group | n | n female | Age (M ± SD) | Dream Recall/week (M ± SD) | LD Frequency Description |
| --- | --- | --- | --- | --- | --- |
| EMS | 14 | 9 | 28.9 ± 6.0 | 4.43 ± 1.18 | 1 none, 8 <1/yr, 4 = 2-3/yr, 1 = 1/mo |
| GVS | 14 | 10 | 26.1 ± 4.2 | 4.07 ± 0.88 | 7 none, 3 <1/yr, 3 = 2-3/yr, 1 = 1/mo |

Note: Demographic information of the participants. LD = lucid dream. EMS = electrical muscle stimulation. GVS = galvanic vestibular stimulation

**Table 2.** Dream Report Overview.

| Group | Total dreams in STIM | Total dreams in SHAM | Final Included Dreams (STIM) | Final Included Dreams (SHAM) | Participants with Dreams in Both | Participants with Only One Condition | Final Participants per group |
| --- | --- | --- | --- | --- | --- | --- | --- |
| EMS | 13 | 16 | 8 | 9 | 4 | 9 | 13 |
| GVS | 12 | 15 | 9 | 10 | 8 | 3 | 11 |

*Note: Reported dreams per participant. Dreams were selected for analysis based on the highest DLQ-7 scores and/or externally rated LD. STIM = stimulation condition, SHAM = sham stimulation condition. EMS = electrical muscle stimulation. GVS = galvanic vestibular stimulation*

### Dream Incorporation

As shown in Table 3, neither EMS, nor GVS stimulation lead to a significantly higher proportion of dreams with direct or indirect incorporation compared to SHAM.

**Table 3.** Dream Incorporation.

| Group | Dream Content | STIM % | SHAM % | $\chi^2$ | p |
| --- | --- | --- | --- | --- | --- |
| EMS (N=17) | Movement of Arms | 37.5% | 33.3% | 0.032 | .858 |
|  | Tactile/Somato Arms | 25% | 0% | 4.26E-06 | .998 |
|  | Direct Incorporation EMS | 37.5% | 0% | 4.70E-04 | .983 |
| GVS (N=19) | Balance Content | 33.3% | 0% | 2.98E-04 | .986 |
|  | Direct Incorporation GVS | 11.1% | 0% | 1.63E-04 | .99 |

*Note: Descriptive results and GLMM analysis of dream incorporation. EMS = Electrical Muscle Stimulation. GVS = Galvanic Vestibular Stimulation.*

### Lucid dreaming

Lucidity was evaluated using externally rated dream reports and DLQ-7 scores (see Table 4).

**Table 4.** GLMM results of Lucid Dreaming Outcomes.

| Group | Measure | STIM | SHAM | $\chi^2/F$ | p |
| --- | --- | --- | --- | --- | --- |
| EMS (N=17) | External Lucidity | 62.5% | 55.6% | $\chi^2 = 0.08$ | .77 |
| | DLQ-7 | M=14.0 ± 10.7 | M=9.3 ± 9.8 | F = 2.46 | 0.175 |
| GVS (N=19) | External Lucidity | 66.7% | 20% | $\chi^2 = 3.84$ | .05* |
| | DLQ-7 | M=14.3 ± 10.8 | M=5.8 ± 4.6 | F = 4.63 | 0.046* |

Note: Descriptive results and GLMM results of externally rated dream reports. DLQ-7 was analysed using a linear mixed model. EMS = electrical muscle stimulation. GVS = galvanic vestibular stimulation. DLQ-7 = Dream Lucidity Questionnaire with item 7 subtracted.

The analysis revealed no significant effect of EMS stimulation condition on externally rated LD or DLQ-7 scores. In the GVS group, subjective lucidity was significantly more frequent in the STIM condition than in SHAM (66.7% vs. 20%, *p* = .05), and DLQ-7 scores differed significantly (p = 0.046).

## Discussion

The present study investigated whether bodily stimulation targeting muscle or vestibular systems can influence lucid dream induction when combined with cognitive training and targeted presleep cueing. Using a 2.5-week sham-controlled laboratory design, we compared EMS and GVS in terms of their effects on dream incorporation and lucidity. Overall, the findings indicate that vestibular stimulation can shape lucid awareness.

### Dream Incorporation and Cue Perception

Consistent with previous work, EMS stimulation was more often incorporated into dream content than SHAM, with direct EMS and somatosensory incorporations observed in 37.5% and 25% of EMS STIM dreams, respectively, whereas no such incorporations occurred in the control condition. However, these findings did not reach significance. This follows previous findings on the incorporation of EMS into dreams (Peters et al., 2024). GVS rarely led to explicit incorporation of the stimulation itself. Balance-related dream content, such as sensations of instability or movement through space, was present in 33.3% of stimulation dreams, compared to none in sham stimulation (p = .986). Similar indirect effects of vestibular stimulation on dream phenomenology have been reported previously, for example, in studies using rocking during sleep or immersive virtual reality before sleep (Leslie & Ogilvie, 1996; Picard-Deland et al., 2020).

### Lucidity Outcomes and Measures

Lucidity was assessed using dream report analysis and questionnaire data, targeting different aspects of the LD experience. EMS stimulation did not significantly increase lucidity relative to control. Importantly, in the EMS condition, lucidity rates were equally high in the control conditions, particularly in the EMS group, compared to previous induction studies (Tan & Fan, 2022). One possible explanation is that lucidity rates were already elevated due to the combination of cognitive training, presleep cueing, and the laboratory context, leaving limited room for additional increases through external stimulation. In addition, the EMS sample contained substantially fewer participants who had never previously experienced a lucid dream compared to the GVS sample. Prior lucid dreaming experience is known to strongly predict induction success (Adventure-Heart, 2020), as individuals with previous lucid dreams might be more likely to recognize unusual dream events and apply cognitive induction strategies effectively. Consequently, the EMS sample may have been more responsive to the cognitive training procedures, leading to relatively high lucidity rates in both stimulation and sham conditions.

In contract, lucidity ratings revealed a significant effect of GVS stimulation, with higher lucidity reports (p = .05) and DLQ-7 scores (p = 0.046) in the stimulation condition compared to the control. Interestingly, rocking participants in a hammock during early-morning REM sleep has been reported to increase lucidity compared to static conditions (Leslie & Ogilvie, 1996), and a presleep virtual reality flight simulation that included full-body vestibular cues increased the likelihood of lucid flying dreams (Picard-Deland et al., 2020). Our use of GVS represents a more targeted approach to vestibular cueing during sleep. Although the stimulation was brief and subtle, the increase in subjective lucidity suggests that vestibular disruptions may influence aspects of dream experience without necessarily being explicitly incorporated into dream narratives. One possible interpretation is that subtle distortions of bodily or spatial processing during dreaming may alter the stability of the dream and linear mixed models for continuous scores), reflecting their different data experience itself. The vestibular system plays a central role in spatial orientation and bodily self-representation (Goldberg et al., 2012; Pfeiffer et al., 2014), and small disruptions to these signals might increase the likelihood that dreamers notice inconsistencies in their experience. Such disruptions could promote reflective awareness without requiring the stimulus itself to appear explicitly in dream content.

### Limitations and Future Directions

Several limitations should be acknowledged. The sample size was small, limiting statistical power and generalizability. Additionally, stimulation parameters, particularly for GVS, were brief (2-second, 1Hz) and subtle, and alternative and optimized timing or intensity parameters may yield different outcomes. The vestibular system typically responds to sustained or dynamic changes in motion (Goldberg et al., 2012), and future studies should explore longer or continuous stimulation to increase incorporation potential (Peters et al., 2024c). EMS occasionally caused premature awakenings, underscoring the need for adaptive stimulation protocols that adjust intensity in real-time to minimize arousals (Pavlou, 2024).

A limitation of the statistical approach is that no formal correction for multiple comparisons (e.g., Bonferroni) was applied. The lucidity outcomes were analysed using different statistical models (GLMM for binary outcomes structures. Although both measures showed a convergent pattern in the GVS condition, the observed effects were close to the conventional significance threshold and based on a modest sample size. The results should therefore be interpreted cautiously. Future research should replicate these findings in larger samples, explore adaptive or prolonged stimulation protocols, and systematically disentangle the contributions of training, expectation, and bodily stimulation. Combining somatosensory and vestibular cues, or tailoring stimulation to individuals, may further clarify how bodily signals can be leveraged to support lucid awareness.

## Conclusion

In summary, the present study suggests that bodily stimulation may influence aspects of dream experience and subjective lucidity in different ways. While EMS occasionally appeared in dreams without increasing lucidity beyond control conditions, GVS was associated with higher subjective lucidity scores despite minimal explicit incorporation into dream reports. These findings suggest that external stimulation during sleep may influence lucid awareness not only through recognizable dream cues, but also through subtle perturbations of bodily and spatial experience. Targeting the dream body may therefore provide a promising avenue for future dream engineering approaches aimed at supporting conscious awareness during sleep.

## Acknowledgments

We would like to extend our gratitude to Susanne Zulauf for helping with the data analysis and to the Technology Platform for Science (TPF) of the Faculty of Human Sciences of the University of Bern.

